# Restriction of chromatin accessibility is necessary for appropriate enhancer expression

**DOI:** 10.1101/311993

**Authors:** Daniel W. Hagey, Susanne Klum, Cecile Zaouter, Jonas Muhr

## Abstract

Tissue specific gene expression underpins cell type diversity, and arises from the cooperative activities of transcription factors and the chromatin landscape. It has been previously demonstrated that enhancers with specific arrangements of transcription factor binding motifs can bring together commonly and specifically expressed factors in order to stabilize chromatin accessibility and drive spatially restricted reporter expression within different regions of the CNS. However, when reporters were used to analyse the activity of enhancers bound differentially by a common factor in the endoderm and CNS, several examples of non-tissue specific reporter expression were observed. In order to judge whether or not this may have been due to the unregulated chromatin environment of exogenously delivered enhancer reporters, here we have analysed the chromatin landscape of cells from the CNS and endodermal tissues and find that this reflects neighbouring gene expression to a greater degree than transcription factor binding. This work demonstrates that chromatin accessibility plays an essential role in defining enhancer activity in distantly related cell types.

---

Cell type specific gene expression is a prerequisite to building complex organisms. In turn, this specificity is dependent on variations between the chromatin landscapes and transcription factor binding profiles of the different cell types (1). It has been demonstrated that specifically expressed transcription factors guide commonly expressed factor’s binding profiles to reflect the gene expression and chromatin landscapes of specific cell type within the same germ layer (2). However, in a subsequent study looking at gene expression specificity between germ layers, reporter expression of several enhancers was found to deviate significantly from that predicted by the transcription factor binding profile (3). One possibility was that enhancer expression specificity using exogenously delivered DNA-constructs was not able to recapitulate the in vivo chromatin accessibility landscape, and thus allowed transcription factor binding where it would not be possible.

In order to assess this possibility, we quantified chromatin accessibility within regions bound by SOX2 in embryonic day (E)11.5 mouse spinal cord, cortex, stomach or lung/esophagus. We took advantage of publically available DNase I-seq data sets produced from the corresponding tissues ((2) and see Material and Methods) and compared them directly by PCA and hierarchical clustering using Diffbind (Figure 1,A and B). Through this, we found that the chromatin accessibility at SOX2 bound regions was much more similar in cell types within the same germ layer than between them.

**Figure 1:**
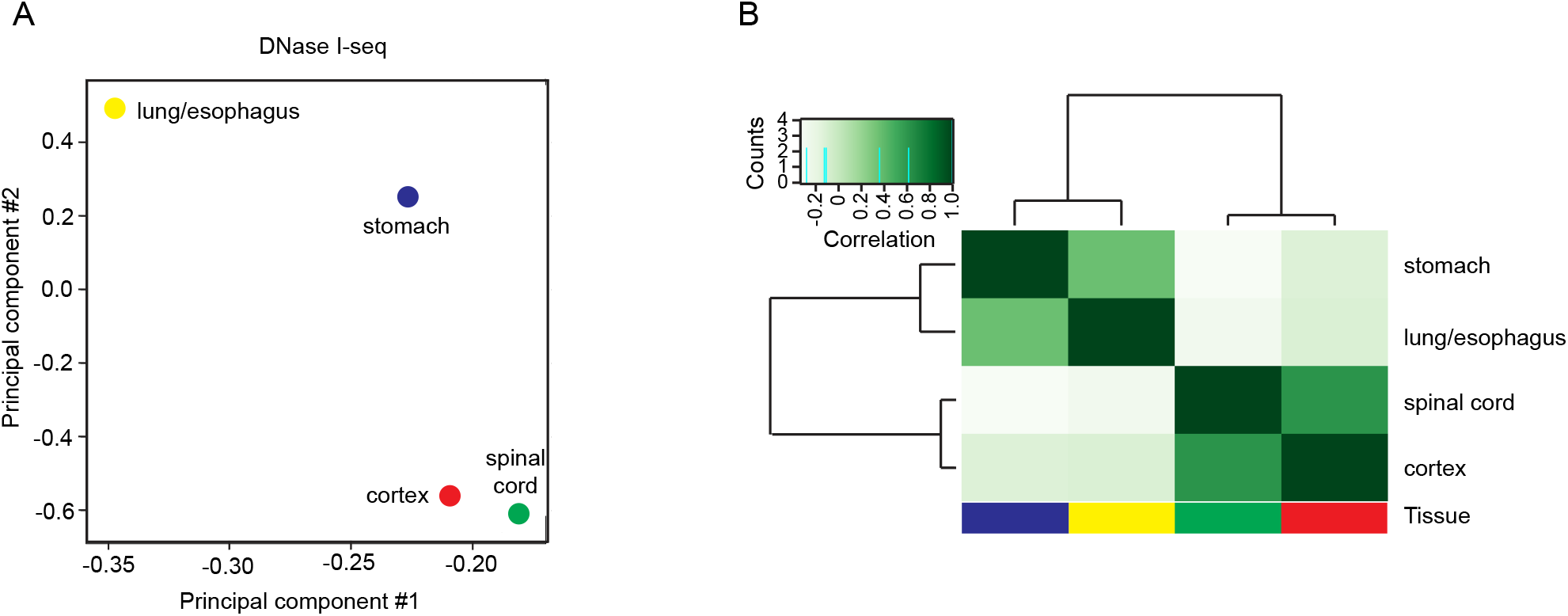
Chromatin accessibility is more similar within a germ layer than between them. (A) Diffbind PCA of the DNase I-seq experiments analysed, based on the read counts in each DNase I-seq at sites of SOX2 ChIP-seq peaks. (B) Diffbind heatmap and dendograms of the correlations between each DNase I-seq experiment analysed; cortex in red, spinal cord in green, stomach in blue and lung/esophagus in yellow.

Next, we could illustrate the divergence in reporter expression patterns generated from stomach specific SOX2 peaks by visualizing Vgll4 +8bp and Foxp1 −10kb enhancer expression in 50hpf embryos (Figure 2A and 2B). To attempt to explain this discrepancy, we separated regions bound by SOX2 specifically in one of the four organs analysed into those that neighboured genes with higher (appropriate tissue expression) or lower (inappropriate tissue expression) than average expression in the tissue that the SOX2 peak was found (Figure 2C). This analysis revealed that SOX2-bound chromatin around genes expressed in a tissue appropriate manner was significantly less accessible in cells of the opposing germ layers than average (Figure 2D). This contrasted with SOX2 targeted enhancers around genes with inappropriate expression profiles in endodermal and neural cells, which were instead found to be accessible in both germ layers (Figure 2D). Hence, the accessibility of DNA-regions bound by SOX2 in a tissue specific manner appears to reflect the expression of associated genes better than the SOX2 binding pattern. Therefore, at least in certain cases, restricted enhancer accessibility appears to be necessary for lineage specific activity, which potentially explains the ectopic activity observed when these enhancers were examined in isolation in transgenic zebrafish embryos.

**Figure 2:**
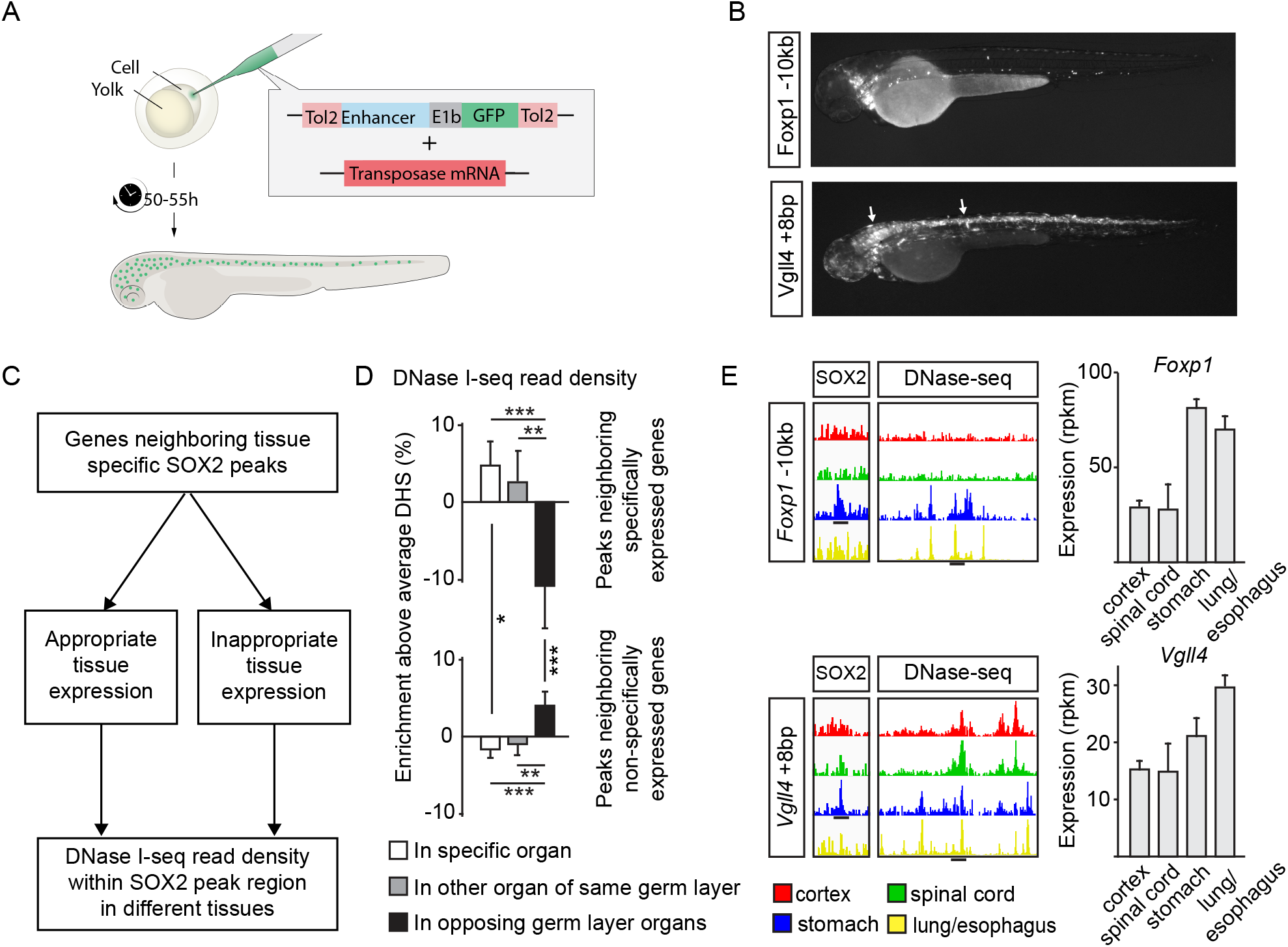
Enhancer chromatin accessibility reflects neighbouring gene expression (A) Schematic of Tol2 based enhancer assay insertion into single cell zebrafish embryos. (B) GFP expression of two stomach specific SOX2-bound enhancers with appropriate (Foxp1 −10kb) and inappropriate (Vgll4 +8bp) reporter patterns in 50hpf zebrafish embryos. White arrows point to CNS reporter expression. (C) Schematic of DNase I-seq read density analysis within cell-type-specific SOX2 ChIP-seq peak regions around appropriately and inappropriately expressed genes. (D) DNase I-seq read depth enrichment over average within cell-type-specific SOX2 bound regions, around appropriately and inappropriately expressed genes in the specific organ (white), the other organ in the same germ layer (grey) and the opposing germ layer (black). Error bars show standard deviation between averages of tissues, and p-values are calculated with two-sided, unpaired t-tests (*=p<0.05, **=p<0.01, ***=p<0.001). (E) Tracks showing SOX2 ChIP-seq and DNase I-seq profiles (cortex in red, spinal cord in green, stomach in blue and lung/esophagus in yellow) and gene expression (RPKMs) for one specifically (*Foxp1*) and one generally (*Vgll4*) expressed gene bound by SOX2 only in the stomach. Black lines under tracks are centred on stomach specific SOX2 peaks.

Recently, much progress has been made in determining the importance of chromatin accessibility and transcription factor activity to gene expression specificity (1–2). Unfortunately, functional assays for the necessity of these inputs have been lacking due to the difficulty of altering the genome and manipulating localized DNA-accessibility in a targeted fashion. We utilized a Tol2-based system to insert enhancer sequences, identified in the mouse genome, upstream of a reporter gene into single cell zebrafish embryos (3). By allowing us to assess the spatial activity of these enhancers outside their native chromatin environment, we establish that restricting chromatin accessibility is necessary for their appropriate activity.

## Ethics statement

All animal procedures and experiments were performed in accordance with Swedish animal welfare laws authorized by the Stockholm Animal Ethics Committee: Dnr N249/14.

### Zebrafish enhancer experiments

Performed according to Hagey et al. 2016

### Diffbind PCAs, dendograms and heatmaps

The PCA plots were created with DiffBind. The plots are based on affinity (read count) data. A binding matrix was calculated with scores based on read counts for every sample within the binding site intervals (peaks). For the DHS PCA plots the peaks of the corresponding Sox2-ChIP were used.

### DNase-seq data and specific gene analysis

DNase-seq data came from Hagey et al. 2016 (cortex and spinal cord) and the ENCODE project (stomach [GEO:GSE83187] and lung [GEO:GSM1014194], both produced by the Stamatoyannopoulous lab). To assess the relationship between DNase-seq read density in regions specifically bound by SOX2 in the different tissues, and the gene expression specificity of neighboring genes, we first assigned all peaks to their nearest gene within 500kb. Any gene expressed below an average of rpkm 1 in FACS sorted Sox2-GFP cells from the four tissues (data from Hagey et al. 2018) was removed from the analysis. Genes were then assigned as being specifically expressed if, for example in genes around cortex specific SOX2 peaks, the proportion of rpkms for a gene in cortex was greater than the overall average rpkm proportion for all genes around cortex specific peaks. In this cortex specific example, the DNase-seq read depth found within the regulatory region in spinal cord corresponds to the value for “In other organ of the same germ layer”, while the read depth found in stomach and lung constitute the value for “In opposing germ layer organs”. These values were then collected, calculated as a ratio to the overall read depth ratio for all specific peaks in cortex, and averaged between all organ and germ layer specificities, with significance assigned using a Student’s t-test to compare these averages.

